# XMAn Update - A Database of *Homo sapiens* Mutated Peptides

**DOI:** 10.64898/2026.08.24.746771

**Authors:** Joshua R. S. Haueis, Iulia M. Lazar

**Affiliations:** Department of Biological Sciences, Virginia Tech, 1981 Kraft Drive, Blacksburg, VA 24061, USA; Fralin Life Sciences Institute/Virginia Tech; Carilion School of Medicine/Virginia Tech

**Keywords:** mass spectrometry, proteogenomics, proteins, database, mutation, SAAVs

## Abstract

Mass spectrometry (MS) is the leading technology for identifying proteins in complex biological samples. It relies on the use of tandem MS alongside a reference database of canonical protein sequences to computationally identify peptides and their parent proteins. The canonical sequences represent the most widely expressed and functionally validated forms of proteins. Consequently, disease-induced or disease-supportive variants, such as those associated with cancer, will evade detection if they are absent from the database. To address this challenge, this study introduces a revised release of the **Unkown M**utation **An**alysis (XMAn) database by incorporating coding missense and nonsense mutations from the latest versions (v103) of the COSMIC Genome Screen Mutants (GSM) and Cancer Gene Census (CGC) datasets in two distinct FASTA-formatted peptide databases comprising 3,848,499 and 312,658 variants, respectively. The mutated peptides were matched to reviewed, non-redundant UniProt *Homo sapiens* protein entries (18,362 and 746), and characterized in terms of nucleotide- and amino acid mutation frequencies, peptide length distributions, and associations between specific single-nucleotide (SNV) and single amino acid (SAAVs) variants. Applied to the analysis of MDA-MB-231 breast cancer cell-membrane protein fractions, the database enabled the identification of 300+ high-quality variant peptides - several localized to functional protein-binding and catalytic domains - and 23 aberrant protein products mapped to the CGC dataset. The database is hosted and available for download on Zenodo (XMAn/gsm doi: 10.5281/zenodo.21781023; XMAn/cgc doi: 10.5281/zenodo.21781514) or can be accessed through https://sites.google.com/vt.edu/xman-db/home.

## Introduction

Proteomic workflows rely on generating alternating mass spectra of parent-peptide and daughter-fragment ions that are interpreted with software packages that map the measured m/z of detected ions to theoretical values and ion patterns generated *in silico* from canonical protein sequences. The core of these software packages consists of embedded search engines that utilize distinct algorithms to score the quality of the mass spectra and identify the amino acid sequence of detected peptides (e.g., cross-correlation-based algorithms such as Sequest, or probability-based algorithms such as Mascot or Andromeda [1], [2], [3], [4]). As the canonical protein sequences from reference databases (DBs) represent the most common, wild-type sequence of a protein, any mutation or modification in the mass of an amino acid will result in a mass shift that will cause the search engine to fail in identifying a match between the experimental and theoretical sequences. Various algorithms that address such limitations have been developed, for example by using *de novo* sequencing [5], or multi-pass error-tolerant and mass-tolerant (open) searches ([6], [7]). The effectiveness of these algorithms continues to be limited, however, by the poor quality or incomplete characteristics of mass spectra in the case of the *de novo* methods, the performance of the first-pass search in the error-tolerant searches, and the high false-discovery rates associated with the mass-tolerant approaches [8]. Novel computational methods incorporate improved versions of previously established algorithms or leverage novel machine learning models to interpret the tandem mass spectra that are missed by conventional search engines ([9], [10], [11], [12], [13], [14]).

In parallel, advances in genomic sequencing technologies have facilitated the completion of massive projects involving thousands to millions of subjects. These efforts have resulted in the generation of extensive datasets that have been curated and compiled in various formats in publicly-available databases, and in the development of sophisticated bioinformatics tools for the processing of such data. **Table 1** summarizes the primary resources for variation analysis including: (i) the major initiatives that provide a comprehensive framework for the integration of genomic, transcriptomic, and proteomic data (COSMIC, GDC/TCGA, PDC and ClinVar [15], [16], [17], [18], [19]); (ii) the platforms that focus on cancer- or cell line-specific analysis (TP53 database, CCLE, DepMap [20], [21], [22], [23]); (iii) the databases that compile inherited disease variants (OMIM, HGMD [24], [25]); and (iv) the results of various efforts aimed at enabling the querying, analysis, and visualization of cancer datasets (cBioPortal, CanProVar, HuVarBase, XMAn, Humsavar [26], [27], [28], [29], [30], [31], [32]). Concurrently, resources with a targeted scope directed at investigating variations in specific protein categories and the functional consequences of molecular alterations (MOKCa-3D, MutHTP, DCMP, ADDRESS [33], [34], [35], [36]), as well as at cataloguing common SNPs (1000 Genomes Project, dbSNP [37], [38]), have been designed. Moreover, recently, a database of artificially generated single nucleotide variants incorporating ∼9 billion entries, along with functional annotations and pathogenicity predictions for the whole human genome, has been developed (VarCards) [39].

**Table 1.**
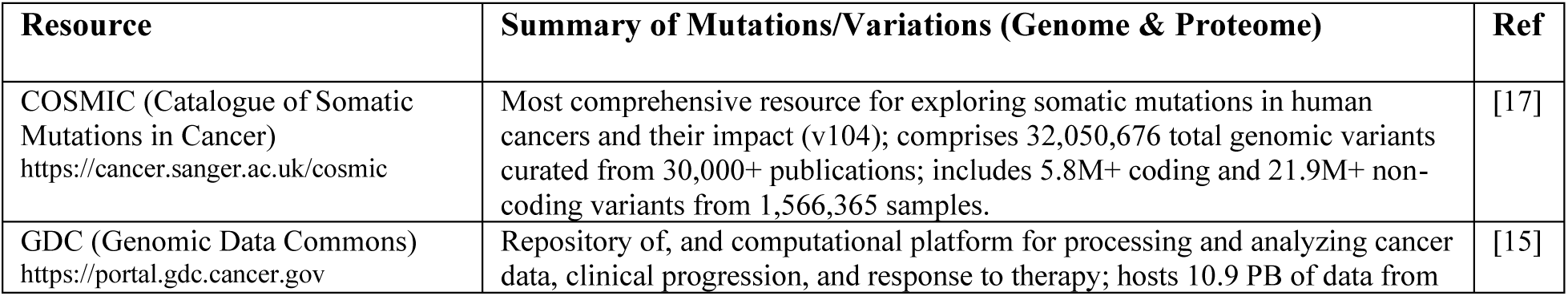
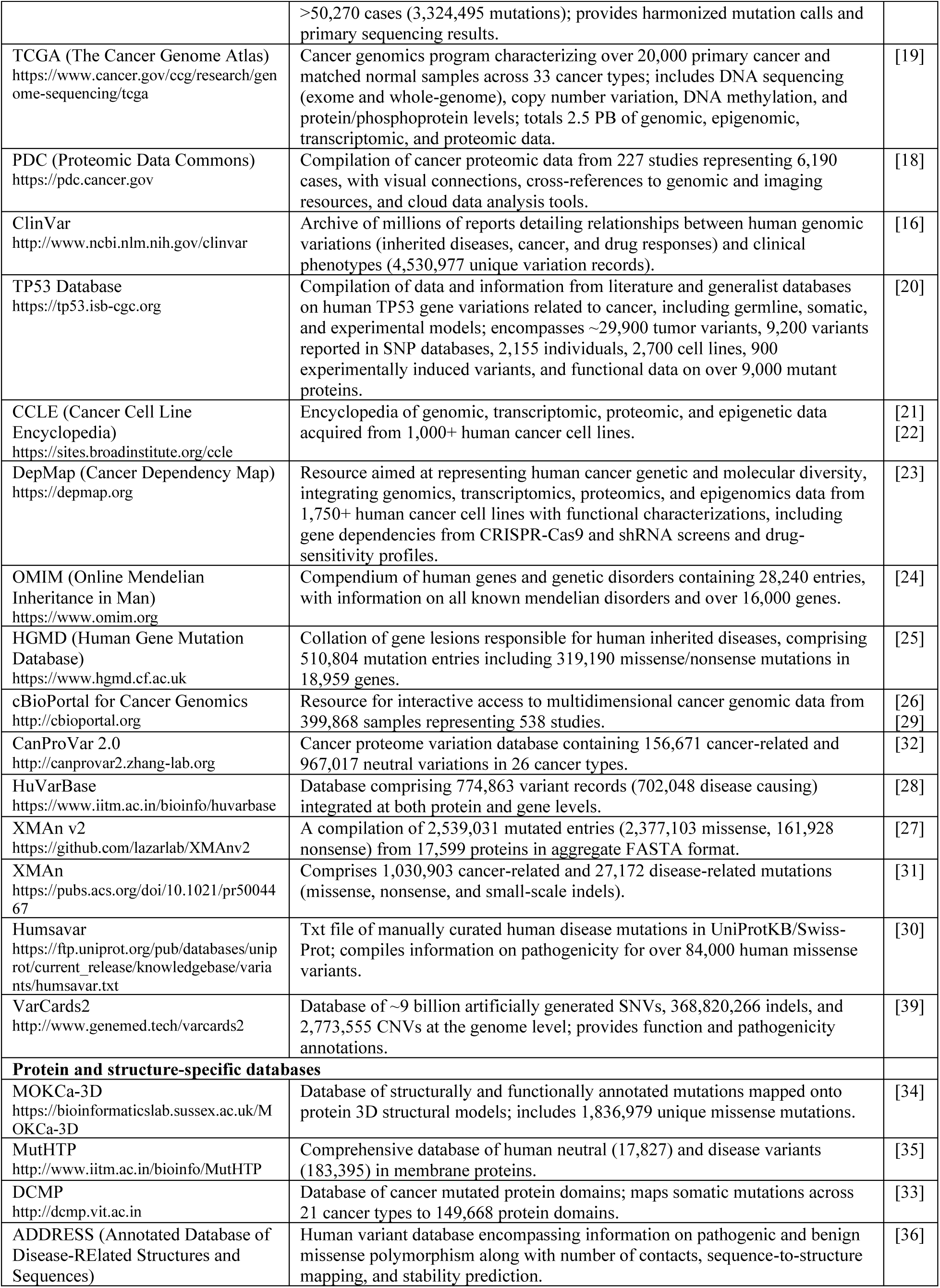
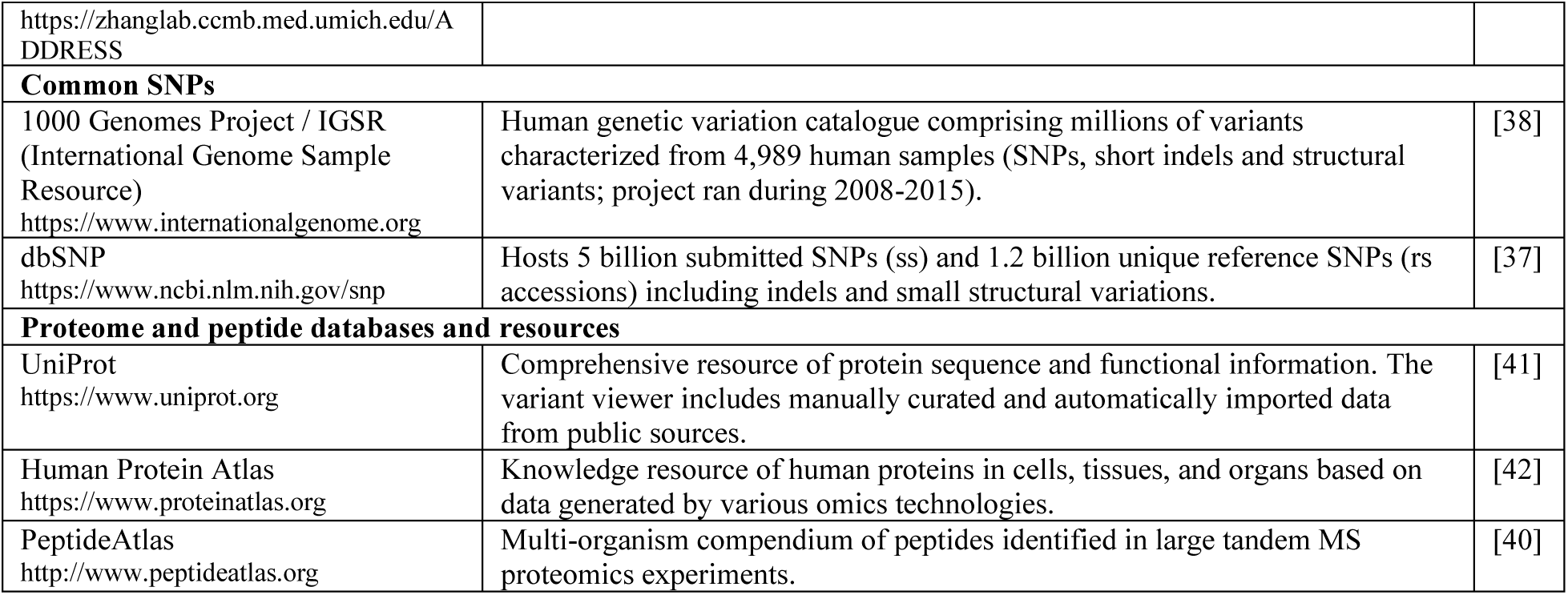
Public resources that archive and functionally characterize human genomic and proteomic variants.

In an effort to complement the large, broad-scope, generalist resources available for proteomics research ([40], [41], [42]), proteogenomics approaches have leveraged the vast archives of millions of documented variants by integrating the genomic and proteomic landscapes through custom-built databases of protein variants tailored to specific categories, individuals, or various physiological states. When appended to the canonical sequences, the combined canonical/mutated databases enable MS search outcomes with improved accuracy and sample-specific identification of protein variants and disease biomarkers. To facilitate the implementation of these strategies, the impact of database inflation on the sensitivity of peptide detection and the underestimation of FDRs has been addressed by developing additional validation and post-search data filtering procedures ([43], [44], [45], [46], [47]). FASTA-formatted, ready-to-use, mutated peptide databases that would allow MS search engines to directly incorporate peptides with altered sequences into their search space without requiring algorithmic modifications are not, however, readily available. To address this gap in proteomics resources and enable the identification of aberrant protein products, in this work, we release an updated version of our previously described XMAn database of mutated peptides ([27], [31]) and provide Python scripting tools for processing the database and proteomic search results. The database was constructed based on the latest release (v103) of the expert-curated Catalogue of Somatic Mutations in Cancer (COSMIC) - the most comprehensive resource for exploring human somatic alterations for identifying cancer driver mutations and discover cancer-specific genetic variants and drug targets. The current version of XMAn integrates the coding missense and non-sense mutations associated with the GSM and the CGC datasets incorporated in two distinct variant peptide databases with 3,848,499 and 312,658 entries, respectively. To the best of our knowledge, to date, this updated version of the XMAn database represents the most extensive compilation of missense and nonsense protein mutations in a FASTA-formatted resource.

## Methods

The variant peptide databases were constructed by using coding nucleotide-level mutation data (point mutations and small indels that induce single amino acid variants) from the COSMIC GSM CGC collections (v103, http://cancer.sanger.ac.uk/cosmic), and full protein amino acid sequences from a reviewed, minimally redundant, manually annotated *Homo sapiens* protein database from UniProt (04/05/2026 download from https://www.uniprot.org, 20,431 entries). A series of in-house developed Python scripts were used to extract information from these datasets and construct the two FASTA files of variant peptides, each encompassing missense and nonsense mutations. In contrast to the previous XMAn versions, the preliminary processing of COSMIC and UniProt data was performed by using the DB Browser for SQLite application (v3.13.1) powered by the lightweight SQLite relational database engine (https://sqlitebrowser.org/dl). This approach allowed for using Structured Query Language (SQL) for streamlining the searching, selection, and retrieval of data imported in the relational databases created from the COSMIC/UniProt downloads.

The variant peptide databases were generated by following a structured workflow. (1) The COSMIC and UniProt (20,431 entries) files were initially imported in the SQLite browser for viewing and determining how to create the best schema for downstream processing of the data. (2) The COSMIC/UniProt SQLite relational databases with relevant attributes were created, deduplicated, and filtered for missense and nonsense mutations. (3) The processed COSMIC and UniProt SQLite files were joined on using gene names as clause. (4) The sequence of mutated peptides was created from the joined COSMIC/UniProt data by a stepwise methodology that included: (i) determining whether the type and position of the mutated amino acid site in the COSMIC DB corresponded to the type and position of the amino acid in the UniProt protein sequence; (ii) determining the sequence of amino acids to the right and left of the mutated site, in case of a positive match; (iii) replacing the original with the mutated amino acid and extracting the left/right sequences including maximum two Lys (K) or Arg (R) residues to either site of the mutation (mimicking, thus, the sequence of tryptic peptides typically generated for proteomic analysis); for nonsense mutations when the nucleotide substitutions generated a terminal codon, only the left sequence to the mutated amino acid was extracted; for variants present near the protein termini, the script extracted only the leftover amino acids in the sequence; for variants where the right/left sequences did not include one or two K/Rs, the flanking sequences were extracted in full [to note that for a few cases that included overlapping patterns at the split boundary such as KKK, KR, etc., the script failed splitting the sequence at the requested boundary - likely due to Python’s string handling process when encountering overlapping patterns; the impact was however minor, resulting only in discrepancies in the number of K/R residues included in the sequence adjacent to the mutation]; (iv) generating a short peptide sequence tag that included the mutated site +/- 3 amino acids to the right and left, for enabling downstream MS filtering of data; and (v) creating a FASTA header in which each mutated peptide entry was identified by a combined UniProt/COSMIC Genomic ID, followed by the COSMIC Legacy mutation ID, the gene and protein names, the protein length, the nucleotide (mutation_CDS) and amino acid mutation types and position in the sequence, the mutated sequence tag, and the descriptions for the mutation, transcript accession, chromosome number, and mutation somatic status (confirmed or of unknown origin). We note that the header captures only the first genomic-level variant from the COSMIC files that was associated with a particular amino acid mutation in a protein, and does not include additional/distinct variants that could have led to identical mutated peptide sequences.

An additional version of the mutated peptide DB (10.5281/zenodo.21856119) was created from COSMIC v100 genome/targeted screen and drug resistance downloads, by using solely Python scripting and the PANDAS package [48]. Supplementary details were included in the header to facilitate biological interpretation [e.g., whether the mutation was part of the protein active/binding site, whether the mutation induced a polarity change, or whether the conversion was between amino acids with identical or close mass that can lead to misassignments (I/L or K/Q)].

MS analysis of the MDA-MB-231 triple negative breast cancer cell-membrane cellular subfractions was performed with a nano EASY-nLC 1200 UHPLC system equipped with a nano-separation column ES802A (250 mm long, 75 μm i.d., 2 μm C18/silica particles, 2 h gradient) interfaced to a Q-Exactive hybrid quadrupole-Orbitrap mass spectrometer (Thermo Fisher Scientific). As the cancer cell-membranes feature an abundant and altered layer of glycoproteins [49], the preparation of the cell-membrane protein fractions was performed by following cell-surface glycoprotein EZ-Link Alkoxyamine-PEG4-Biotin labeling/NeutrAvidin pulldown protocols. The cell-membrane protein enrichment method and the LC-MS/MS data dependent acquisition parameters were described in detail in prior work [50]. A set of 18 raw MS files were generated from three biological replicates of serum-deprived and serum-rich MDA-MB-231 cell cultures, analyzed in triplicate by LC-MS/MS.

Data analysis was performed with the Proteome Discoverer 3.1 software package and the Sequest HT search engine. The search was performed with a three-pass workflow that used the Percolator peptide spectrum match (PSM) validator node to discriminate between correct and incorrect PSMs. In the 1^st^ and 2nd passes, the DB search was performed against the *Homo sapiens* database only, PSMs with FDR>1 being passed on to the next search. In the 3^rd^ pass, the DB search was performed against the mutated peptide database. All throughout, the Percolator parameters were set to concatenated target/decoy validation based on q-value, maximum Delta Cn 0.05, and FDR stringent/relaxed targets of 0.01/0.05. The Sequest node search parameters included fully tryptic precursor peptides with maximum 2 missed cleavages, min/max 6/50 amino acids within a range of 400-5000 m/z, b/y and neutral loss ions included only, and mass tolerances set to 10 ppm for the precursor and 0.02 Da for the fragment ions. Dynamic modifications were pass-specific, and were chosen to maximize the number of matches to the canonical database in the first two passes, as follows: (i) 1^st^ pass: peptide oxidation (+15.995 Da on M); peptide terminus none; protein N-terminus acetylation (+42.011 Da on N-terminus), Met loss (-131.040 Da on M), and Met loss+acetylation (-89.030 Da on M); (ii) 2^nd^ pass: peptide deamidation (+0.984 Da on N/Q); peptide terminus Gln > pyro-Glu (-17.027 Da on Q), carbamylation (+43.006 Da on N-terminus); protein N-terminus acetylation (+42.011 Da on N-terminus), Met loss (-131.040 Da on M), and Met loss+acetylation (-89.030 Da on M); (iii) 3^rd^ pass (only the most commonly observed modifications were applied): peptide oxidation (+15.995 Da on M) and deamidation (+0.984 Da on N/Q). For the final report, only unique variant peptide sequences with Rank 1, Xcorr≥1.5, Sequest node PEP≤0.01, and with no additional match to any canonical UniProt ID, were selected. Isobaric I>L or L>I conversions were removed from the Proteome Discoverer output report. Lastly, a Python script was developed to discard any peptide sequences with “Master Protein Accession” identifier that did not contain the actual mutation site. This was necessary because the mutated peptides contain two K/R residues on both sides of the mutation, which enables the reporting of peptide segments that exclude the actual amino acid variant. These can be eliminated by selecting only sequences that contain the full 7 amino acid long peptide variant tag, or at least the first or last 4 amino acids of the tag provided in the FASTA header.

Supplementary Python scripts were developed for processing the information in the mutated database and can be downloaded from https://github.com/lazarlab/Proteomics_db_tools. The scripts include code for (i) counting the number of entries or unique sequences in the XMAn database, (ii) re-writing the DB with selections of mutated sequences within a min-max range of amino acids, (iii) creating gene-specific derivative databases of variant peptides, and (iv) removing deceptive peptide matches from the proteomics search results exported to Excel that are annotated with UniProt/COSMIC ID but do not include the actual mutated site. Google/Gemini and OpenAI/ChatGPT were used in some instances to assist with troubleshooting and building the Python scripts. Stacked bar charts and histograms were generated with Excel and circular dendograms with RawGraphs. Dendograms were edited with the Boxy SVG editor. Venn diagrams were built with tools provided by https://www.biotools.fr/. SIFT and PolyPhen-2 scores were extracted either from uniprot.org or calculated with tools provided by PolyPhen-2 (https://genetics.bwh.harvard.edu/pph2/bgi.shtml).

## Results

SQLite-based processing of the COSMIC files streamlined the accurate selection of the desired mutations and elimination of duplicate entries, the present XMAn update nearly doubling the entry counts of the most recent release [27]. With consistent COSMIC revisions, these numbers are expected and likely to continue increase. Relevant examples of peptide-level missense and nonsense mutations derived from single-nucleotide variants and small indels - as displayed in the XMAn FASTA files - are provided in **Table 2**, and the overall statistics for the GSM and CGC mutated peptide files are captured in **Table 3**. Analysis of the data revealed distinct differences in the frequency of missense vs. nonsense mutations, reflecting the functional impact of these alterations across the genome. **Figures 1A-C** display the specific SNVs and SAAVs extracted from the GSM dataset. As many of the CGC peptides overlapped with the GSM peptides (>76 %), the trends for the CGC dataset closely mirrored those of the GSM set.

**Figure 1.**
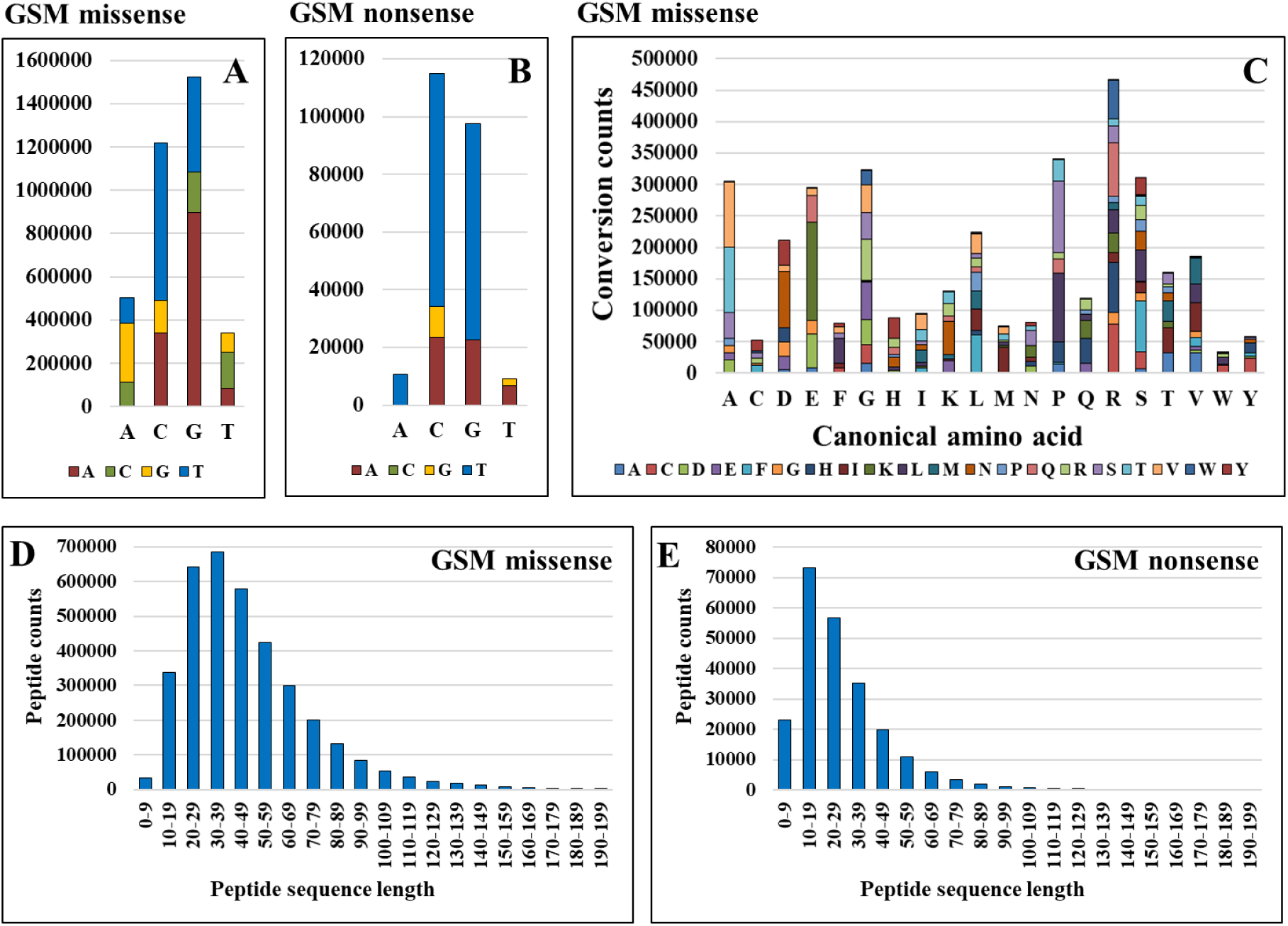
Characterization of the XMAn database in terms of nucleotide- and amino acid mutation frequencies (COSMIC/GSM dataset). (**A**)/(**B**) SNV missense and nonsense counts; (**C**) SAAV missense counts; (**D**)/(**E**) Histograms representing the distribution of missense and nonsense peptide lengths (x-axis truncated at a length of 200 amino acids).

**Table 2.**
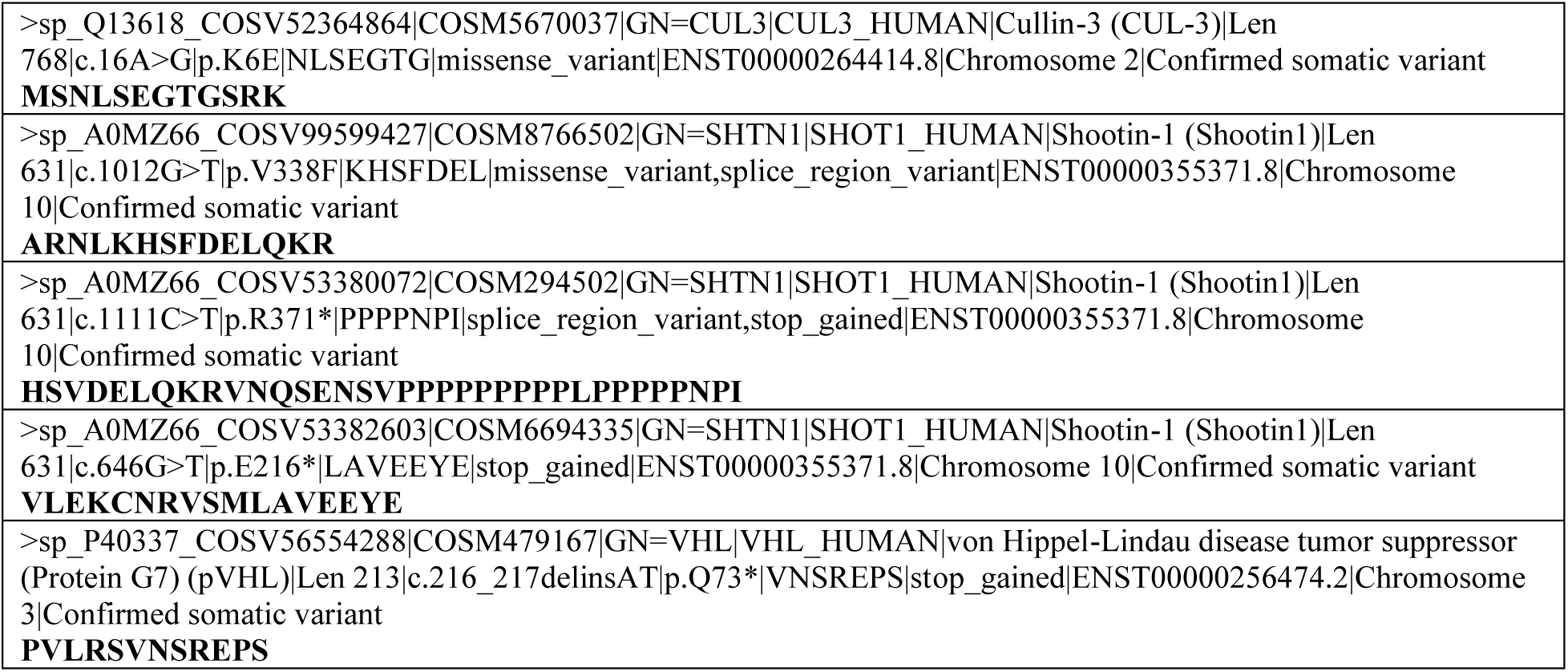
Somatic and nonsense peptide entries from the XMAn database that exemplify the FASTA entry heading structure.

**Table 3.** Characterization of the mutated lists of peptides in the XMAn database subsets.

|  | **GSM (COSMIC v103)** |  | **CGC (COSMIC v103)** |  |
| --- | --- | --- | --- | --- |
|  | Missense | Nonsense | Missense | Nonsense |
| Unique peptides | 3,613,786 | 234,713 | 286,954 | 25,704 |
| Minimum length | 6 | 6 | 6 | 6 |
| Maximum length | 1,680 | 1,015 | 1,680 | 1,015 |
| Mean length | 49 | 28 | 58 | 31 |
| SD | 42 | 23 | 91 | 33 |
| Median length | 41 | 23 | 43 | 24 |

The largest number of missense mutations were generated by cytosine (C) to thymine (T) and guanine (G) to adenine (A) conversions (**Figure 1A**). These represent coding mutations, identifying the actual molecular changes at the transcript level (not genomic level, for which the “-“ strand codes the opposite nucleotide pair). The data reflect the chemical vulnerability of the methylated cytosines at CpG dinucleotide sites, where the highly unstable 5-methylcytosines spontaneously lose an amine group and convert into a thymine. While unmethylated CpG islands are generally protected from this deamination mechanism, methylated CpGs outside these islands drive the highest rates of spontaneous point mutations in the human genome. The perceived high “frequency” of recorded G>A mutations is not however due to G instability. Aside of processes that induce G>A alterations (e.g., DNA replication errors such as misincorporation or mispairing, DNA repair errors, oxidative DNA damage), this conversion rather mirrors the primary C>T mutation captured on the complementary DNA strand after replication establishes a permanent T:A base pair (i.e., the DNA replication process replaces the C:G with the T:A pair). For nonsense mutations (**Figure 1B**), the nucleotide alterations reflected only changes that generated termination codons that couldn’t have been created by conversions to new C bases (i.e., TAA, TAG, and TGA).

The effect of nucleotide-level changes on the relative frequency of amino acid mutations is provided in **Figure 1C**, with the greatest impact being experienced by the A, D, E, G, L, P, R, and S amino acids, specifically, in terms of A>T, A>V, D>N, E>K, G>R, L>F, P>L, P>S, R>H, R>Q, R>W, and S>F conversions. The largest number of mutations were accumulated by R (i.e., losses in R), even when normalized to the frequency of amino acids in the human proteome [31]. This is not unexpected, as arginine is encoded by 6 codons of which 4 contain a CpG dinucleotide [51]. Conversions to K, L, S, and V were the largest. Analysis of variant peptides lengths exposed that the median values in the mutated databases were ∼41 for the missense and ∼23 for the nonsense populations (**Table 3**), and that most of the peptide sequences contained <120 amino acids (**Figure 1D,E**). This range provides MS detection opportunities for up to 50-60 amino acid long peptides flanking the mutation site, falling within the conventional 5,000 Da mass limit utilized by database search engines, regardless of missed K/R cleavages. Gains and losses in K, R, P, D, and E are expected however to affect the tandem mass spectra quality and sequence assignments. To increase protein identification stringency, the database can be truncated to a specific sequence length range, for example 9-120 amino acids (see available Python scripts in the methods section).

The correlation between nucleotide and amino acid mutations revealed that the nucleotide C>T and G>A conversions were the main drivers of the high mutation frequencies observed in the above-highlighted amino acids (A, D, E, G, L, P, R, and S) (**Figure 2**). At their specific site, these non-conservative mutations alter the amino acid polarity (N, Q, S, T), acid/base properties (D, E, H, K, R), or bulkiness (F, P, W), thereby affecting overall protein structure, conformation, function, and even stability/half-life. Many such alterations functionally impair key tumor suppressor and DNA damage repair proteins. For example, R175H, R248Q, R248W, R273H, and R282W are critical hotspot mutations within the DNA-binding domain of the p53 tumor suppressor, a master regulator of essential cellular processes such as cell cycle arrest, senescence, apoptosis, and genomic stability. Together with R249S, R273S, and G245S, these variants account for ∼28 % of all cancer-associated p53 mutations [52], with the p53 protein presenting a disrupted function in over 50 % of all human cancers.

**Figure 2.**
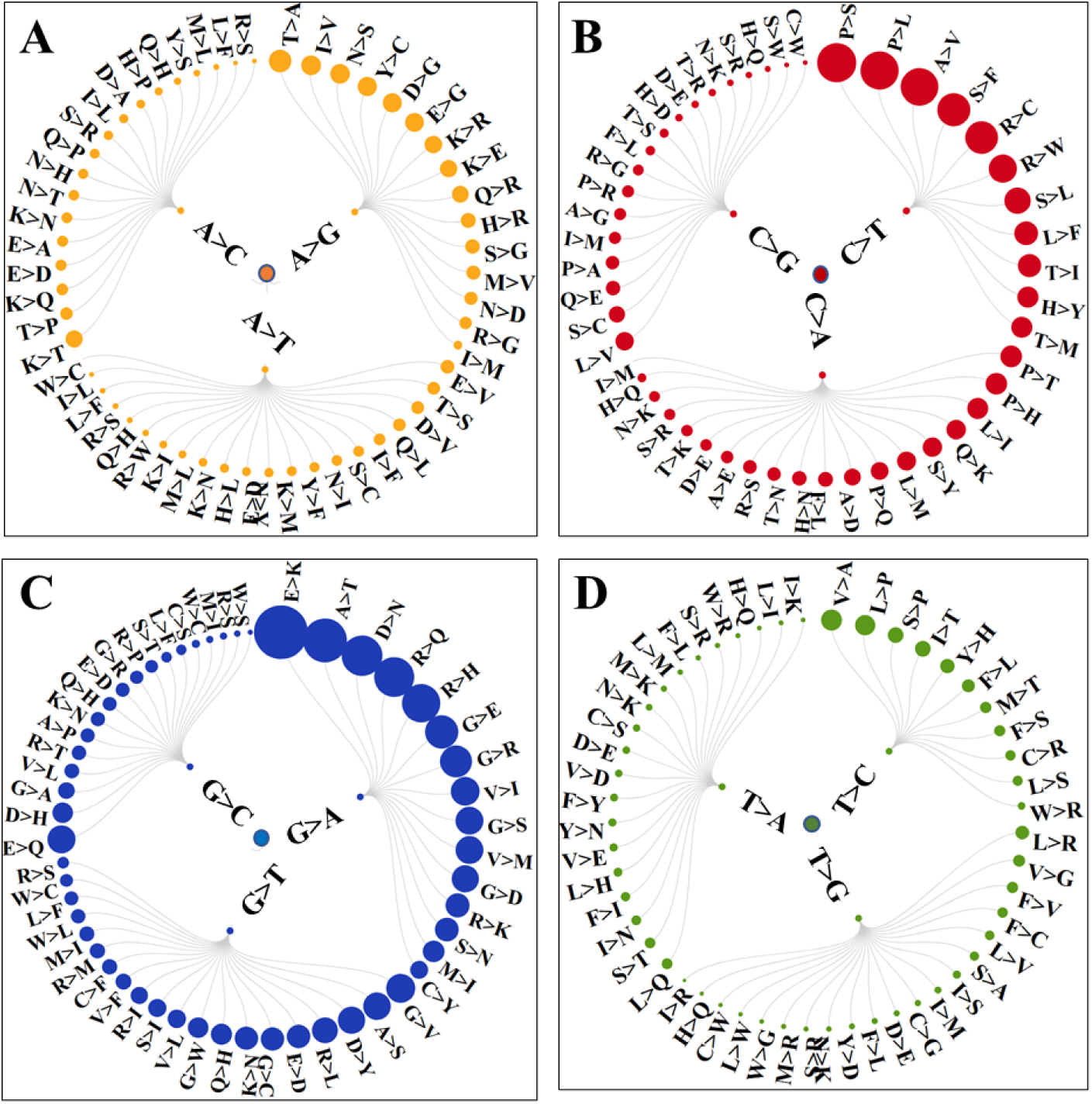
Dendograms depicting the associations between specific SNV and SAAVs in the XMAn database. SNVs were grouped based on the originating nucleotide in the canonical sequence: (A) adenine; (B) cytosine; (C) guanine; (D) thymine. SAAV bullets sizes are proportional to the number of recorded mutants.

Protein-level validation of genomic variants is not straightforward and often unsuccessful. Genomic diversity and instability can lead to multiple and confounding experimental outcomes. The genetic code is degenerate, therefore, not all SNVs will result in altered proteins sequences, and synonymous SNVs will not affect the amino acid sequence of a protein either. The ability to detect SAAVs is impacted by protein half-life, posttranslational modifications, low or altered protein expression, co-expression of both alleles (50 % wild-type/50 % mutant), and quick clearing of misfolded proteins or pathogenic variants by the cellular machinery before detection can be achieved. These challenges can be further compounded by an altered enzymatic cleavage pattern that leads to incomplete sequence coverage by MS detection.

The value of integrating the XMAn database in the proteomic data analysis workflow was evaluated by analyzing the cell-membrane fraction of the MDA-MB-231 triple negative breast cancer cell line. Using our established protocols [50], we identified ∼1,900 cell-membrane and cell-periphery associated proteins in these cells (to be discussed in future work). Raw data reprocessing by using the UniProt/XMAn GSM and UniProt/CGC databases resulted in the identification of 314 and 41 mutated peptide sequences (29 overlaps) matched to 219 and 19 human proteins, respectively, several carrying multiple mutated sites. **Table 4** provides a comparative analysis of results, demonstrating that a proper analysis strategy and choice of database search parameters can effectively handle the extended search space of variant peptides.

**Table 4.**
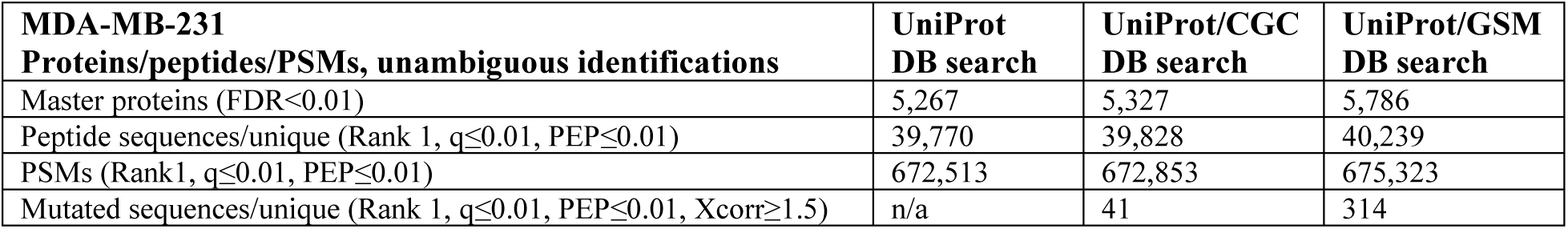
Comparative analysis of database search results by using UniProt and UniProt/variant databases.

| **MDA-MB-231**<br>**Proteins/peptides/PSMs, unambiguous identifications** | **UniProt**<br>**DB search** | **UniProt/CGC**<br>**DB search** | **UniProt/GSM**<br>**DB search** |
| --- | --- | --- | --- |
| Master proteins (FDR<0.01) | 5,267 | 5,327 | 5,786 |
| Peptide sequences/unique (Rank 1, $q \leq 0.01$, $PEP \leq 0.01$) | 39,770 | 39,828 | 40,239 |
| PSMs (Rank1, $q \leq 0.01$, $PEP \leq 0.01$) | 672,513 | 672,853 | 675,323 |
| Mutated sequences/unique (Rank 1, $q \leq 0.01$, $PEP \leq 0.01$, $Xcorr \geq 1.5$) | n/a | 41 | 314 |

The identification of canonical peptides was not affected by the use of variant databases in the workflow, all canonical sequences being identifiable in the subsequent multi-database searches (**Figure 3A**). Per need, a stringency-based tiering approach can be used to increase the number and confidence of identified variants, for example by customizing mass tolerance (Δm), using canonic/variant-specific FDRs, or using isolation interference filters. Here, mass accuracy measurements were ≤5 ppm for ∼80 % and ≤2 ppm for ∼60 % of the mutated sequences. Filtering the PSMs by % isolation interference can effectively isolate very high-quality tandem mass spectra, but requires careful implementation to mitigate the risk of discarding valid identifications. The data processing pipeline can be further refined for targeted analysis of small subsets of variants (see methods for the availability of Python scripts). A large, readily-available, curated variant database with FASTA-patterned headers such as XMAn enables users to create a broad range of custom datasets by simply filtering the headers for specific keywords. For example, a dedicated tumor protein p53 derivative file can thus be instantly generated by filtering the CGC database for the keyword "GN=TP53.” Using this approach we extracted a list of 1,432 TP53-specific mutated peptides, provided as a FASTA file in **Supplemental file 1**. Similarly, a cancer-specific signature database can be constructed by inputting a list of multiple gene names.

**Figure 3.**
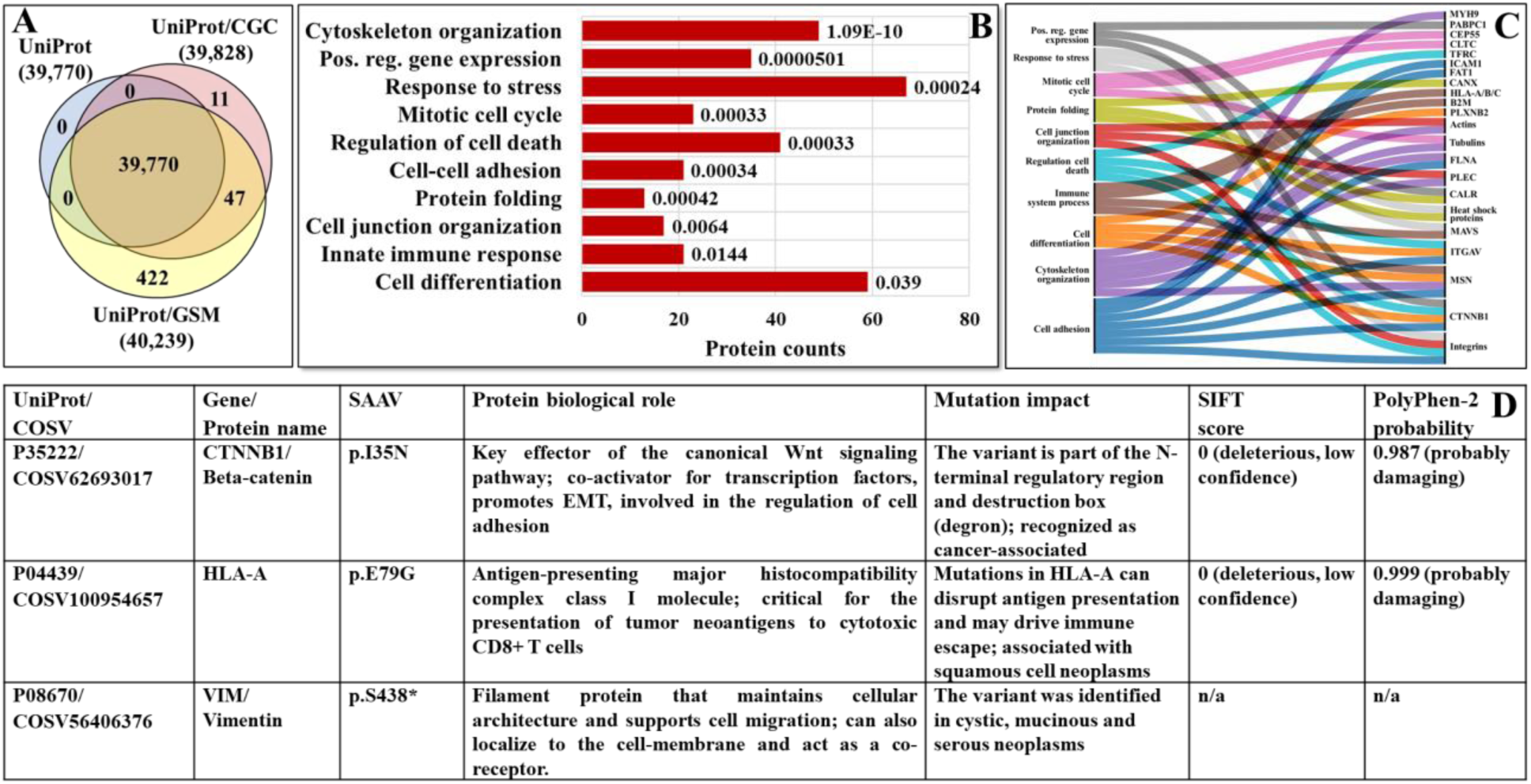
Cancer-supportive biological processes and proteins for which missense and nonsense mutations were identified. (**A**) Venn diagram of protein identifications by using UniProt or UniProt/mutated databases. (**B**) Selection of GO biological processes to which the identified protein variants could be mapped (the labels represent enrichment FDRs). (**C**) Alluvial diagram linking relevant examples of mutated proteins (or protein categories) to biological processes affected in cancer. (**D**) Examples of protein variants with possible association to cancer.

While a comprehensive discussion of the extended biological impact of the detected SAAVs is beyond the scope of this study, several key observations warrant highlighting. GO classification of the variant proteins resulted in multiple enriched biological categories of relevance to cancer progression (**Figure 3B**). Along with several cell-membrane proteins (CGC database), it was not surprising to observe many mutated actin-, tubulin-, integrin-, and heat shock proteins (GSM database) – which are stable, highly abundant proteins in the cell. A few non-cell-membrane proteins, likely contaminants from the cytosolic/nuclear, cytoskeleton-associated, or mitochondrial cellular subfractions were also present (e.g., PKM, MAVS, CEP55, ERCC2). The cell-membrane localization of certain nuclear proteins, e.g., DNA repair proteins, in metastatic tumors has been however previously reported, and can be used as an additional indicator of pathogenicity (https://atlasgeneticsoncology.org). Such proteins have additional non-repair roles in cell signaling or binding to ECM components, or can act as (co-)receptors. While a large number of the observed mutations were single-nucleotide polymorphisms, many clustered within the protein binding or catalytic domains and have yet unknown functional impact on disease. For example, mutations that disrupt the cell cycle and cell differentiation processes can lead to uncontrolled proliferation, while mutations that alter the mechanisms of cellular metabolism, cell death/apoptotic processes, or immune regulation, can further drive malignant transformation. Altogether, 23 mutated proteins could be mapped to the cancer-driver genes included in the CGC dataset (ATIC, ATP1A1, B2M, CALR, CLTC, CTNNB1, ERCC2, FAT1, FLNA, FUS, HLA-A, ITGAV, LASP1, MSN, MYH9, NONO, NPM1, PABPC1, PLEC, RAP1B, SDHA, TFRC, TPM4), which together with the GSM-derived variants underscored the capabilities enabled by the use of the XMAn database in the proteomic workflow (**Figure 3C**). Lastly, mutations that were previously associated with cancer were also detected in the MDA-MB-231 cells. Three examples are provided in **Figure 3D**. The pathogenicity SIFT (deleterious if <0.05) and PolyPhen (damaging if >0.9) scores predict whether an amino acid substitution will affect protein structure and function. While not cancer-driver mutations, for products validated by gene-level alterations, the cumulative multi-protein variant burden can serve as an indicator of functional impact and disease liability.

## Conclusions

With over 3.8 million variant peptide entries, the updated XMAn FASTA files enable protein-level confirmation of genomic variants and the identification of a wide range of aberrant protein products. As the mutated sequences can be appended to standard reference databases, and search parameters and post-filtering strategies can be optimized without altering the underlying code of established search engines, the identification of variant peptides is straightforward. The inclusion of short sequence snippets in the FASTA headers enables successful elimination of false variant matches. The distinct COSMIC/CGC-based peptide database enables focused proteomic searches concentrated on Cancer Gene Census entries only. Custom-tailored derivative databases can be easily created from the main FASTA files by selecting gene-specific mutation subsets. Practical implementation in a proteomic workflow resulted in the identification of a large number of protein variants implicated in a broad spectrum of cancer-supportive biological processes. In future studies, the ready-to-use, aggregate format of the XMAn files will serve as an effective platform for onco-proteogenomics discoveries. The integration of genome-level alterations with proteome-level structural and functional outcomes into freely accessible resources will help assess the cumulative impact of somatic mutations, accelerate the discovery of mutational signatures of disease, and advance the design of effective therapies.

## Supporting information

FASTA-formatted subset of TP53 mutated peptides extracted from the XMAn database

## Acknowledgements

This work was supported in part by an award from the National Institute of General Medical Sciences (R01-GM121920) to IML. Mutation datasets were downloaded from the Sanger Institute Catalogue of Somatic Mutations in Cancer, releases v103 and v100 (http://cancer.sanger.ac.uk/cosmic). The research conception, design, and interpretation of results are the sole intellectual contribution of investigators.

## Author Contributions

IML and JH developed the Python scripts, built the database, and wrote the paper. IML coordinated the work.

## References

[1] J. Cox, N. Neuhauser, A. Michalski, R. A. Scheltema, J. V. Olsen, and M. Mann, “Andromeda: A Peptide Search Engine Integrated into the MaxQuant Environment,” J. Proteome Res., vol. 10, no. 4, pp. 1794–1805, Apr. 2011, doi: 10.1021/pr101065j.

[2] J. K. Eng, A. L. McCormack, and J. R. Yates, “An approach to correlate tandem mass spectral data of peptides with amino acid sequences in a protein database,” J. Am. Soc. Mass Spectrom., vol. 5, no. 11, pp. 976–989, Nov. 1994, doi: 10.1016/1044-0305(94)80016-2.

[3] A. I. Nesvizhskii et al., “Dynamic Spectrum Quality Assessment and Iterative Computational Analysis of Shotgun Proteomic Data,” Molecular & Cellular Proteomics, vol. 5, no. 4, pp. 652–670, Apr. 2006, doi: 10.1074/mcp.M500319-MCP200.

[4] D. N. Perkins, D. J. C. Pappin, D. M. Creasy, and J. S. Cottrell, “Probability-based protein identification by searching sequence databases using mass spectrometry data,” Electrophoresis, vol. 20, no. 18, pp. 3551–3567, Dec. 1999, doi: 10.1002/(SICI)1522-2683(19991201)20:18<3551::AID-ELPS3551>3.0.CO;2-2.

[5] C. C. A. Ng, Y. Zhou, and Z.-P. Yao, “Algorithms for de-novo sequencing of peptides by tandem mass spectrometry: A review,” Analytica Chimica Acta, vol. 1268, p. 341330, Aug. 2023, doi: 10.1016/j.aca.2023.341330.

[6] J. M. Chick et al., “A mass-tolerant database search identifies a large proportion of unassigned spectra in shotgun proteomics as modified peptides,” Nat Biotechnol, vol. 33, no. 7, pp. 743–749, Jul. 2015, doi: 10.1038/nbt.3267.

[7] D. M. Creasy and J. S. Cottrell, “Error tolerant searching of uninterpreted tandem mass spectrometry data,” Proteomics, vol. 2, no. 10, pp. 1426–1434, Oct. 2002, doi: 10.1002/1615-9861(200210)2:10<1426::AID-PROT1426>3.0.CO;2-5.

[8] R. Salz et al., “Personalized Proteome: Comparing Proteogenomics and Open Variant Search Approaches for Single Amino Acid Variant Detection,” J. Proteome Res., vol. 20, no. 6, pp. 3353–3364, Jun. 2021, doi: 10.1021/acs.jproteome.1c00264.

[9] S. Choi and E. Paek, “MutCombinator: identification of mutated peptides allowing combinatorial mutations using nucleotide-based graph search,” Bioinformatics, vol. 36, no. Supplement_1, pp. i203–i209, Jul. 2020, doi: 10.1093/bioinformatics/btaa504.

[10] K. Karunratanakul, H.-Y. Tang, D. W. Speicher, E. Chuangsuwanich, and S. Sriswasdi, “Uncovering Thousands of New Peptides with Sequence-Mask-Search Hybrid De Novo Peptide Sequencing Framework,” Molecular & Cellular Proteomics, vol. 18, no. 12, pp. 2478–2491, Dec. 2019, doi: 10.1074/mcp.TIR119.001656.

[11] C. Landerer, M. Scheremetjew, H. Moon, L. Hersemann, and A. Toth-Petroczy, “deTELpy: Python package for high-throughput detection of amino acid substitutions in mass spectrometry datasets,” Bioinformatics, vol. 40, no. 7, p. btae424, Jul. 2024, doi: 10.1093/bioinformatics/btae424.

[12] S. Wang, M. Zhu, and B. Ma, “NeoMS: Mass Spectrometry-Based Method for Uncovering Mutated MHC-I Neoantigens,” IEEE Trans. Comput. Biol. Bioinform., vol. 22, no. 2, pp. 444–454, Mar. 2025, doi: 10.1109/TCBB.2024.3447746.

[13] M. Yilmaz et al., “Sequence-to-sequence translation from mass spectra to peptides with a transformer model,” Nat Commun, vol. 15, no. 1, p. 6427, Jul. 2024, doi: 10.1038/s41467-024-49731-x.

[14] C. Zhu et al., “Identification of non-canonical peptides with moPepGen,” Nat Biotechnol, vol. 44, no. 4, pp. 568–573, Apr. 2026, doi: 10.1038/s41587-025-02701-0.

[15] A. P. Heath et al., “The NCI Genomic Data Commons,” Nat Genet, vol. 53, no. 3, pp. 257– 262, Mar. 2021, doi: 10.1038/s41588-021-00791-5.

[16] M. J. Landrum et al., “ClinVar: public archive of relationships among sequence variation and human phenotype,” Nucl. Acids Res., vol. 42, no. D1, pp. D980–D985, Jan. 2014, doi: 10.1093/nar/gkt1113.

[17] Z. Sondka et al., “COSMIC: a curated database of somatic variants and clinical data for cancer,” Nucleic Acids Research, vol. 52, no. D1, pp. D1210–D1217, Jan. 2024, doi: 10.1093/nar/gkad986.

[18] R. R. Thangudu et al., “NCI’s Proteomic Data Commons: A Cloud-Based Proteomics Repository Empowering Comprehensive Cancer Analysis through Cross-Referencing with Genomic and Imaging Data,” Cancer Research Communications, vol. 4, no. 9, pp. 2480–2488, Sep. 2024, doi: 10.1158/2767-9764.CRC-24-0243.

[19] The Cancer Genome Atlas Research Network et al., “The Cancer Genome Atlas Pan-Cancer analysis project,” Nat Genet, vol. 45, no. 10, pp. 1113–1120, Oct. 2013, doi: 10.1038/ng.2764.

[20] K. C. De Andrade et al., “The TP53 Database: transition from the International Agency for Research on Cancer to the US National Cancer Institute,” Cell Death Differ, vol. 29, no. 5, pp. 1071–1073, May 2022, doi: 10.1038/s41418-022-00976-3.

[21] M. Ghandi et al., “Next-generation characterization of the Cancer Cell Line Encyclopedia,” Nature, vol. 569, no. 7757, pp. 503–508, May 2019, doi: 10.1038/s41586-019-1186-3.

[22] Y. Li et al., “Proteogenomic data and resources for pan-cancer analysis,” Cancer Cell, vol. 41, no. 8, pp. 1397–1406, Aug. 2023, doi: 10.1016/j.ccell.2023.06.009.

[23] A. Tsherniak et al., “Defining a Cancer Dependency Map,” Cell, vol. 170, no. 3, pp. 564–576.e16, Jul. 2017, doi: 10.1016/j.cell.2017.06.010.

[24] A. Hamosh, “Online Mendelian Inheritance in Man (OMIM), a knowledgebase of human genes and genetic disorders,” Nucleic Acids Research, vol. 33, no. Database issue, pp. D514–D517, Dec. 2004, doi: 10.1093/nar/gki033.

[25] P. D. Stenson et al., “The Human Gene Mutation Database (HGMD®): optimizing its use in a clinical diagnostic or research setting,” Hum Genet, vol. 139, no. 10, pp. 1197–1207, Oct. 2020, doi: 10.1007/s00439-020-02199-3.

[26] E. Cerami et al., “The cBio Cancer Genomics Portal: An Open Platform for Exploring Multidimensional Cancer Genomics Data,” Cancer Discovery, vol. 2, no. 5, pp. 401–404, May 2012, doi: 10.1158/2159-8290.CD-12-0095.

[27] M. A. Flores and I. M. Lazar, “XMAn v2—a database of *Homo sapiens* mutated peptides,” Bioinformatics, vol. 36, no. 4, pp. 1311–1313, Feb. 2020, doi: 10.1093/bioinformatics/btz693.

[28] K. Ganesan, A. Kulandaisamy, S. Binny Priya, and M. M. Gromiha, “HuVarBase: A human variant database with comprehensive information at gene and protein levels,” PLoS ONE, vol. 14, no. 1, p. e0210475, Jan. 2019, doi: 10.1371/journal.pone.0210475.

[29] J. Gao et al., “Integrative Analysis of Complex Cancer Genomics and Clinical Profiles Using the cBioPortal,” Sci. Signal., vol. 6, no. 269, Apr. 2013, doi: 10.1126/scisignal.2004088.

[30] The UniProt Consortium, “UniProt: a hub for protein information,” Nucleic Acids Research, vol. 43, no. D1, pp. D204–D212, Jan. 2015, doi: 10.1093/nar/gku989.

[31] X. Yang and I. M. Lazar, “XMAn: A *Homo sapiens* Mutated-Peptide Database for the MS Analysis of Cancerous Cell States,” J. Proteome Res., vol. 13, no. 12, pp. 5486–5495, Dec. 2014, doi: 10.1021/pr5004467.

[32] M. Zhang et al., “CanProVar 2.0: An Updated Database of Human Cancer Proteome Variation,” J. Proteome Res., vol. 16, no. 2, pp. 421–432, Feb. 2017, doi: 10.1021/acs.jproteome.6b00505.

[33] I. A. Emerson and K. K. Chitluri, “DCMP: database of cancer mutant protein domains,” Database, vol. 2021, p. baab066, Apr. 2022, doi: 10.1093/database/baab066.

[34] B. Haile et al., “MOKCa-3D database: functional and structural analysis of missense mutations in cancer,” Database, vol. 2026, p. baag001, Jan. 2026, doi: 10.1093/database/baag001.

[35] A. Kulandaisamy et al., “MutHTP: mutations in human transmembrane proteins,” Bioinformatics, vol. 34, no. 13, pp. 2325–2326, Jul. 2018, doi: 10.1093/bioinformatics/bty054.

[36] J. Woodard, C. Zhang, and Y. Zhang, “ADDRESS: A Database of Disease-associated Human Variants Incorporating Protein Structure and Folding Stabilities,” Journal of Molecular Biology, vol. 433, no. 11, p. 166840, May 2021, doi: 10.1016/j.jmb.2021.166840.

[37] L. Phan et al., “The evolution of dbSNP: 25 years of impact in genomic research,” Nucleic Acids Research, vol. 53, no. D1, pp. D925–D931, Jan. 2025, doi: 10.1093/nar/gkae977.

[38] The 1000 Genomes Project Consortium et al., “A global reference for human genetic variation,” Nature, vol. 526, no. 7571, pp. 68–74, Oct. 2015, doi: 10.1038/nature15393.

[39] Z. Wang et al., “VarCards2: an integrated genetic and clinical database for ACMG-AMP variant-interpretation guidelines in the human whole genome,” Nucleic Acids Research, vol. 52, no. D1, pp. D1478–D1489, Jan. 2024, doi: 10.1093/nar/gkad1061.

[40] F. Desiere, “The PeptideAtlas project,” Nucleic Acids Research, vol. 34, no. 90001, pp. D655–D658, Jan. 2006, doi: 10.1093/nar/gkj040.

[41] The UniProt Consortium et al., “UniProt: the Universal Protein Knowledgebase in 2025,” Nucleic Acids Research, vol. 53, no. D1, pp. D609–D617, Jan. 2025, doi: 10.1093/nar/gkae1010.

[42] M. Uhlen et al., “Towards a knowledge-based Human Protein Atlas,” Nat Biotechnol, vol. 28, no. 12, pp. 1248–1250, Dec. 2010, doi: 10.1038/nbt1210-1248.

[43] S. Aggarwal, A. Raj, D. Kumar, D. Dash, and A. K. Yadav, “False discovery rate: the Achilles’ heel of proteogenomics,” Briefings in Bioinformatics, vol. 23, no. 5, p. bbac163, Sep. 2022, doi: 10.1093/bib/bbac163.

[44] J. A. Alfaro, A. Ignatchenko, V. Ignatchenko, A. Sinha, P. C. Boutros, and T. Kislinger, “Detecting protein variants by mass spectrometry: a comprehensive study in cancer cell-lines,” Genome Med, vol. 9, no. 1, p. 62, Dec. 2017, doi: 10.1186/s13073-017-0454-9.

[45] P. Blakeley, I. M. Overton, and S. J. Hubbard, “Addressing Statistical Biases in Nucleotide-Derived Protein Databases for Proteogenomic Search Strategies,” J. Proteome Res., vol. 11, no. 11, pp. 5221–5234, Nov. 2012, doi: 10.1021/pr300411q.

[46] H. Li, Y. S. Joh, H. Kim, E. Paek, S.-W. Lee, and K.-B. Hwang, “Evaluating the effect of database inflation in proteogenomic search on sensitive and reliable peptide identification,” BMC Genomics, vol. 17, no. S13, p. 1031, Dec. 2016, doi: 10.1186/s12864-016-3327-5.

[47] S. Verbruggen et al., “Spectral Prediction Features as a Solution for the Search Space Size Problem in Proteogenomics,” Molecular & Cellular Proteomics, vol. 20, p. 100076, 2021, doi: 10.1016/j.mcpro.2021.100076.

[48] J. R. Haueis, “A Mutated Peptide Database for the Analysis of Proteomic Mass Spectrometry Data.” 2025. [Online]. Available: https://vtechworks.lib.vt.edu/items/440d2c33-4888-4900-9c02-abdd623e0aca.

[49] I. M. Lazar, J. Deng, F. Ikenishi, and A. C. Lazar, “Exploring the glycoproteomics landscape with advanced MS technologies,” Electrophoresis, vol. 36, no. 1, pp. 225–237, Jan. 2015, doi: 10.1002/elps.201400400.

[50] A. Karcini and I. M. Lazar, “The SKBR3 cell-membrane proteome reveals telltales of aberrant cancer cell proliferation and targets for precision medicine applications,” Sci Rep, vol. 12, no. 1, p. 10847, Jun. 2022, doi: 10.1038/s41598-022-14418-0.

[51] J.-C. Walser and A. V. Furano, “The mutational spectrum of non-CpG DNA varies with CpG content,” Genome Res., vol. 20, no. 7, pp. 875–882, Jul. 2010, doi: 10.1101/gr.103283.109.

[52] E. H. Baugh, H. Ke, A. J. Levine, R. A. Bonneau, and C. S. Chan, “Why are there hotspot mutations in the TP53 gene in human cancers?,” Cell Death Differ, vol. 25, no. 1, pp. 154–160, Jan. 2018, doi: 10.1038/cdd.2017.180.

